# A *Ralstonia solanacearum* type III effector alters the actin and microtubule cytoskeleton to promote bacterial virulence in plants

**DOI:** 10.1101/2023.11.01.565113

**Authors:** Rachel Hiles, Abigail Rogers, Namrata Jaiswal, Weiwei Zhang, Jules Butchacas, Marcus V. Merfa, Taylor Klass, Erica Kaser, Jonathan M. Jacobs, Christopher J. Staiger, Matthew Helm, Anjali S. Iyer-Pascuzzi

## Abstract

Cellular responses to biotic stress frequently involve signaling pathways that are conserved across eukaryotes. These pathways include the cytoskeleton, a proteinaceous network that senses external cues at the cell surface and signals to interior cellular components. During biotic stress, dynamic cytoskeletal rearrangements serve as a platform from which early immune-associated processes are organized and activated. Bacterial pathogens of plants and animals use proteins called type III effectors (T3Es) to interfere with host immune signaling, thereby promoting virulence. We previously found that RipU, a T3E from the soilborne phytobacterial pathogen *Ralstonia solanacearum* K60 (*Rs* K60), co-localizes with the plant cytoskeleton. Here, we show that RipU from *Rs* K60 (RipU^K60^) physically associates with both actin and tubulin and disrupts actin and microtubule cytoskeleton organization. We find that pharmacological disruption of the tomato (*Solanum lycopersicum*) cytoskeleton promotes *Rs* K60 colonization. RipU^K60^ suppresses cell surface-triggered immune responses including flg22-mediated reactive oxygen species (ROS) production and callose deposition. Importantly, tomato plants inoculated with *Rs* K60 lacking RipU^K60^ (Δ*ripU^K60^*) had reduced wilting symptoms and significantly reduced root colonization when compared to plants inoculated with wild-type *Rs* K60. Collectively, our data suggest that *Rs* K60 uses the type III effector RipU^K60^ to remodel cytoskeletal organization, thereby promoting pathogen virulence.

## Introduction

Microbial pathogens use virulence proteins known as type III effector proteins (T3Es) to cause disease in plants and animals. T3Es are secreted and retained in the plant cell extracellular space (apoplast) or translocated into the plant cell cytoplasm where they subsequently interact with and manipulate host proteins to suppress a diverse range of processes, including immune signaling [1–3]. The cytoskeleton is an intracellular filamentous network that is essential for cellular homeostasis and is a critical part of immune signaling in eukaryotes [4–7]. Several T3Es from plant and animal pathogenic bacteria interact directly or indirectly with components of the cytoskeleton and suppress immune responses [1,7–10]. The underlying cellular mechanisms for how pathogen-derived effectors manipulate the cytoskeleton, how such manipulation interferes with immune signaling, and how this promotes pathogen virulence remain largely unknown. Knowledge as to how pathogenic bacteria manipulate cellular targets will likely provide insight into host-microbe cellular biology as well as contribute to our understanding of putative disease control strategies [11,12].

*Ralstonia solanacearum* (*Rs*) is a soil-borne pathogen that causes bacterial wilt disease in over 250 plant species, including economically and agriculturally important cash crops such as potato (*Solanum tuberosum*), pepper (*Capsicum annum*), and tomato (*Solanum lycopersicum*) [13–17]. Similar to other pathogenic bacteria, *Rs* utilizes T3E proteins to promote its virulence, but host targets are not well defined [18,19]. In most crop species, host genetic resistance to *Rs* is a quantitative trait and relies upon the action of several genomic regions known as quantitative trait loci (QTL) [17,20–22]. Despite the importance of *Rs* worldwide, mechanisms underlying this quantitative resistance remain largely unknown, but likely involve developmental and basal immune processes [13,16,23].

Immune signaling pathways and their associated proteins are frequent targets of T3Es in plants and animals. During early invasion by microbes, eukaryotic host cells recognize Microbe Associated Molecular Patterns (MAMPs). MAMP recognition elicits a set of downstream signaling events, including production of reactive oxygen species (ROS), calcium (Ca^2+^) influx, and Mitogen Activated Protein Kinase (MAPK) phosphorylation, which ultimately lead to changes in host defense gene transcription and together are known as pattern-triggered immunity (PTI) [24]. The cytoskeleton is an essential signaling intermediate during PTI [4,5,7]. This intracellular filamentous network controls cell shape and division, organellar movement, endocytosis and secretion, and provides the channels for intracellular and extracellular trafficking. In plants, the cytoskeleton is composed of two major filament systems, actin and microtubules, along with many additional accessory proteins that are required for cytoskeletal function [25]. During immune signaling in both plants and animals, the actin cytoskeleton is dynamically linked to ROS burst and Ca^2+^ changes, promote antimicrobial protein transport, and facilitate immune receptor dynamics at the plasma membrane [4,5,7,26–28]. In both plants and animals, inhibiting cytoskeleton dynamics with either genetic mutants or pharmacological inhibitors can promote pathogen virulence [27,29–31].

Both actin and microtubules transiently polymerize and depolymerize in response to internal and external stimuli, including beneficial and pathogenic microbes [4,5,7]. For example, fungal and oomycete invasion promotes actin filament accumulation at the attempted point of penetration [32–35]. Additionally, a transient increase in the density of cortical actin filaments occurs within minutes after bacterial MAMP perception in Arabidopsis (*Arabidopsis thaliana*) and is an early marker of PTI [29,36,37]. Arabidopsis mutants defective in early actin remodeling or dynamics are more susceptible to the bacterial pathogen *Pseudomonas syringae* pv. *tomato* DC3000 (*Pst* DC3000) [29] and have altered immune outputs including callose production and defense gene activation [36–40]. Significant changes in microtubule organization in response to MAMPs or pathogen perception are less well characterized. Microtubule reorganization occurs in response to both beneficial fungi and during pathogen invasion [41,42]. How MAMP recognition leads to changes in cytoskeleton organization and dynamics, and how this remodeling promotes immune signaling and resistance remains largely unknown, although several actin binding proteins are required for changes in actin organization [36,37,40].

The underlying cellular mechanisms for how pathogen-derived effectors manipulate the cytoskeleton, how such manipulation interferes with immune signaling, and how this promotes pathogen virulence remain largely unknown. Nevertheless, several T3Es from phytopathogenic bacteria target either the actin or microtubule cytoskeleton. For example, the *Pseudomonas syringae* effectors HopW1 and HopG1 [30,43,44] as well as XopR from *Xanthomonas campestris* [39] alter actin structure and organization either by directly interacting with actin filaments (HopW1) or by interfering with actin associated proteins (XopR and HopG1). Additional T3Es impact microtubules either directly or indirectly. XopL from *X. euvesicatoria* directly interacts with microtubules and causes cell death when transiently expressed in *N. benthamiana* [45]. Transient expression of HopZa1 from *P. syringae* causes destruction of microtubule networks, inhibits protein secretion, and suppresses cell wall-mediated defenses [31]. The T3Es HopE1 and AvrBsT indirectly impact microtubule organization by interfering with microtubule associated proteins MAP65 [46] and ACIP1 [47], respectively.

We previously showed that RipU^K60^, a T3E from *Rs* strain K60, qualitatively co-localizes with the actin cytoskeleton in tomato roots and leaves of *N. benthamiana*, suggesting it may have a functional role in cytoskeleton-mediated immune signaling [19]. Here, using high-resolution spinning disk confocal microscopy (SDCM) and quantitative image analysis, we demonstrate that RipU^K60^ alters the organization of both the actin and microtubule cytoskeleton. We find that RipU^K60^ physically associates with both tubulin and actin as well as suppresses host immune responses including reactive oxygen species production and callose deposition in response to the MAMP elicitor, flagellin22 (flg22). Cytoskeleton disruption using the pharmacological inhibitors LatrunculinB (LatB; actin) or oryzalin (microtubules) promotes *Rs* colonization in tomato roots, demonstrating that the cytoskeleton has a functional role in *Rs* recognition. A *Rs* mutant lacking RipU^K60^ has decreased virulence and colonization in naturalistic soil drench assays. Collectively, our data suggest that RipU^K60^ promotes *Rs* virulence likely by associating and interfering with the dynamics and organization of both the actin and microtubule cytoskeleton.

## Results

### A Rs Type III effector co-localizes with the cytoskeleton

Using laser scanning confocal microscopy we previously showed that the *Rs* K60 T3E RipU^K60^ qualitatively co-localized with actin [19]. To assess this association quantitatively and extend the analysis to the microtubule cytoskeleton, we investigated the subcellular localization of RipU^K60^- GFP using spinning disk confocal microscopy (SDCM) following transient co-expression in *N. benthamiana* leaves with cytoskeletal reporters. After confirming that RipU^K60^ is secreted through the type III secretion system (Supplemental Figure 1), we imaged and quantified the subcellular co-localization of RipU^K60^-GFP with either the actin reporter fABD2-mCherry or the microtubule reporter TUB5-mCherry. RipU^K60^ and the fluorescent reporters were transiently expressed in *N. benthamiana* leaves using *Agrobacterium tumefaciens*-mediated transient transformation and imaged using SDCM at 48-hours post-infiltration (hpi). At 48 hpi, RipU^K60^ modestly (Pearsons correlation coefficient (Pcc) = 0.3) but significantly co-localized with fABD2-mCherry and prominently (Pcc = 0.6) with TUB5-mCherry (Figure 1). Another *Rs* K60 T3E, RipBD-GFP, that localizes to the plasma membrane serves as a negative control for these experiments and did not demonstrate significant (Pcc values < 0.15) co-localization with microtubules or actin filaments (Figure 1). Thus, RipU^K60^ co-localizes with both actin and tubulin in *N. benthamiana* epidermal cells.

**Figure 1.**
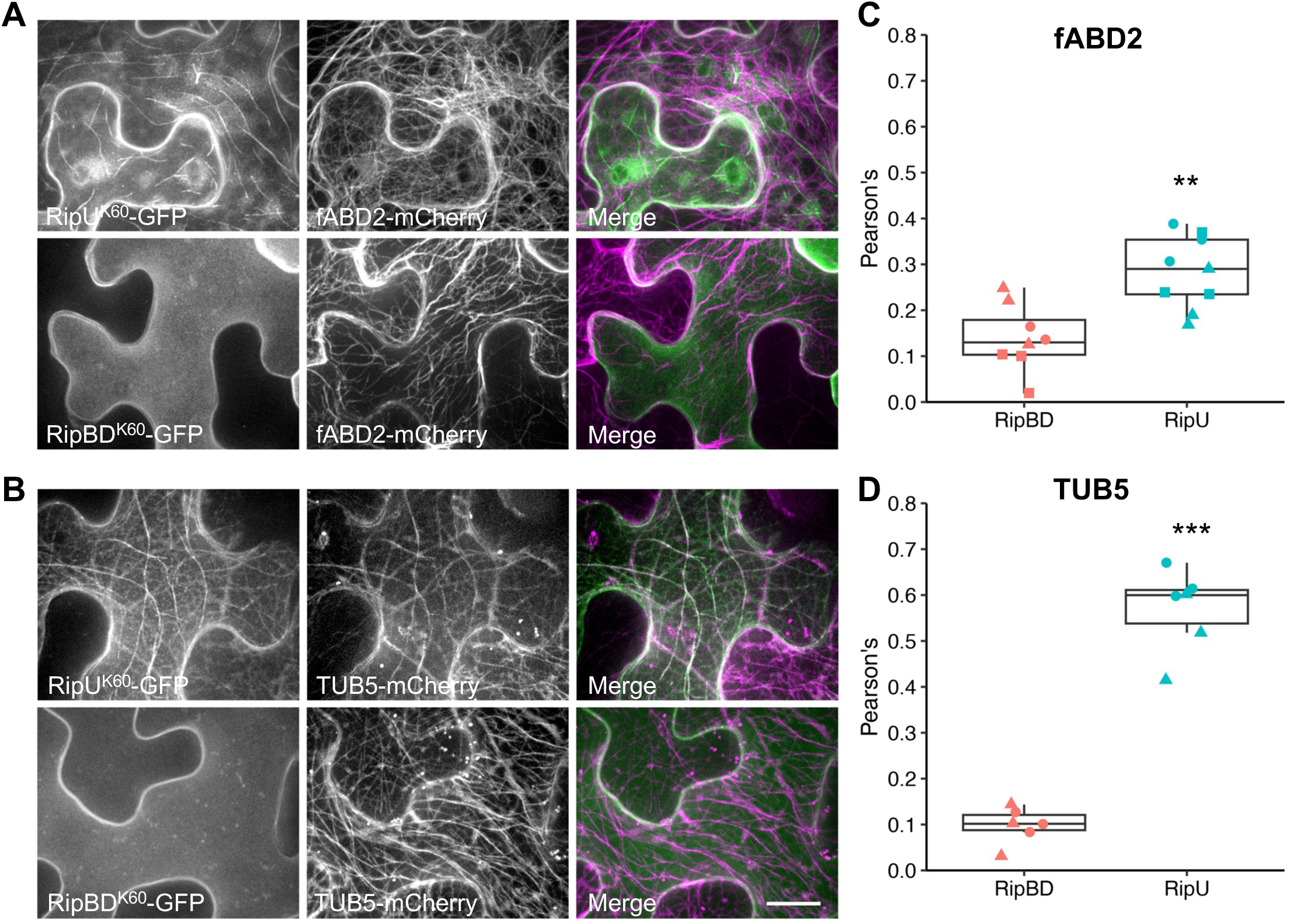
RipU^K60^ colocalizes with the plant cytoskeleton. RipU^K60^-GFP or RipBD^K60^-GFP were co-infiltrated with the actin marker fABD2-mCherry (A) or the microtubule reporter TUB5-mCherry (B) in *N. benthamiana* leaves. Scale bar = 20 µm. Co-localization was quantified at 48 hours post infiltration using Pearson’s coefficient analysis (C, D). RipU^K60^ significantly co-localizes with both actin and microtubules compared to RipBD^K60^ (C, F). 5-15 cells were measured at each infiltration site and the values were averaged as one biological sample; n = 3 for each biological repeat; Data points with different shapes represent different biological repeats; T test; ** P < 0.01; *** P < 0.001.

### RipU^K60^ physically associates with actin and tubulin

The observation that RipU^K60^ co-localized with both actin and microtubule reporters suggested that this T3E protein may physically interact with the components of the plant cytoskeleton. To investigate whether RipU^K60^ associates with actin and tubulin, we transiently expressed RipU^K60^- GFP and performed coimmunoprecipitation (co-IP) assays. Both actin and tubulin were detected in the RipU^K60^-GFP immunoprecipitates (Figure 2). As a control, we transiently expressed free GFP, which did not immunoprecipitate actin or tubulin (Figure 2). These data suggest that RipU^K60^ physically associates with components of the plant cytoskeleton.

**Figure 2.**
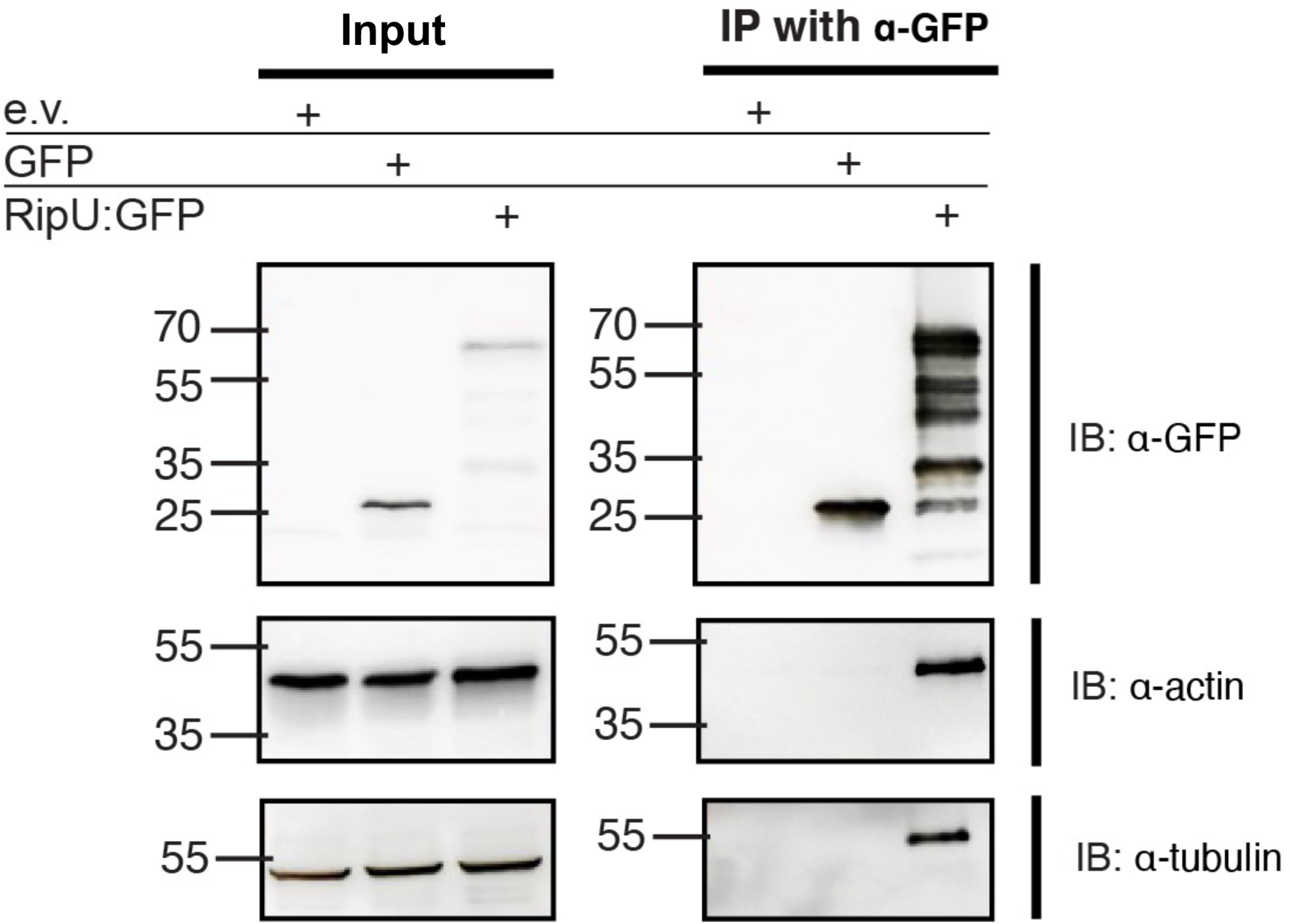
RipU^K60^-GFP co-immunoprecipitates with actin and tubulin. The indicated constructs were transiently expressed in *N. benthamiana* leaves. All transgenes were under the control of a 35S promoter. Total protein was isolated 48 hpi, immunoprecipitated by GFP-Trap agarose bead slurry, and immunoblotted with the indicated antibodies.

### RipU^K60^ suppresses cell surface-triggered immune responses but not programmed cell death

Having shown that RipU^K60^ localizes to and physically associates with components of the plant cytoskeleton, we next sought to gain insight into the putative function of this effector by testing whether RipU^K60^ could suppress reactive oxygen species (ROS) production and callose deposition mediated by flagellin22 (flg22) from *Pseudomonas aeruginosa*. Transient expression of RipU^K60^-GFP in *N. benthamiana* leaves consistently suppressed flg22-mediated ROS production when compared to the empty vector control (Figure 3A). We next investigated whether RipU^K60^ could suppress callose deposition. We generated stable transgenic *Arabidopsis* lines that express HA-tagged RipU^K60^ under the control of a dexamethasone-inducible promoter (pDEX-RipU^K60^-HA). Given that *Rs* is a root invading pathogen, we examined the impact of RipU^K60^ protein expression on flg22*^Pa^*-elicited callose production in root tissues. Qualitative analysis showed that RipU^K60^ expression suppressed flg22*^Pa^*-induced callose deposition in transgenic Arabidopsis roots (Figure 3B).

**Figure 3.**
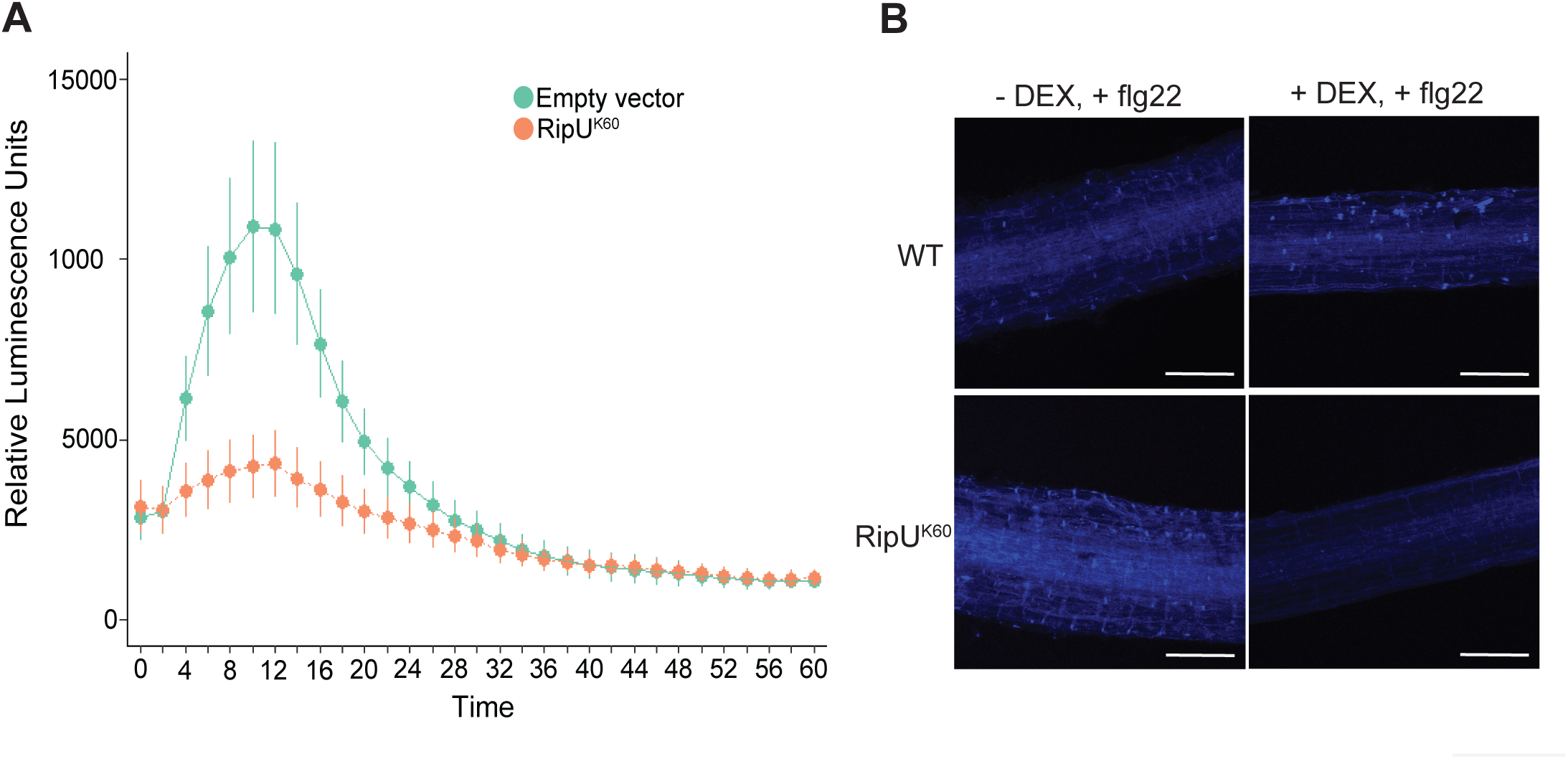
RipU^K60^ suppresses cell surface-triggered immune responses. *N. benthamiana* leaves were infiltrated with either pB7FWG2-RipU^K60^-GFP or pB7FWG2 empty vector and challenged with flg22. (A) flg22-elicited reactive oxygen species production was reduced in pB6FWG2-RipU^K60^-GFP treated leaves when compared to the empty vector treatment. (B) Dexamethasone induced expression of RipU^K60^ in *A. thaliana* suppresses flg22-elicited callose deposition. Scale bar is 100 µm.

In addition to suppressing flg22*^Pa^*-mediated cell surface-triggered immune responses, several *Rs* T3Es suppress hypersensitive response (HR)-like cell death [18,48]. Hence, we investigated whether RipU^K60^ could suppress defense-related cell death triggered by the mammalian pro-apoptotic protein Bcl-2 associated X protein (BAX). To this end, we transiently expressed RipU^K60^-GFP in *N. benthamiana* and, twenty-four hours later, agroinfiltrated BAX and assessed cell death suppression. Intriguingly, RipU^K60^-GFP was unable to suppress BAX-mediated cell death, suggesting that this effector is not likely to function as a general suppressor of cell death (Supplemental Figure 2).

### RipU^K60^ contributes to pathogenesis and virulence

Given our findings that RipU^K60^ contributes to host immune suppression we hypothesized that RipU^K60^ is required for *Rs* K60 virulence and pathogenicity. To test this, we generated a Δ*ripU^K60^* single mutant and Δ*ripU* miniTn*7*::*ripU*K60 (hereafter Δ*ripU^K60^::RipU^K60^*), the complemented *Rs* K60 strain. We first compared colonization rates of wild-type *Rs* K60, the Δ*ripU^K60^* single mutant, and the Δ*ripU^K60^::RipU^K60^*complemented strain in resistant and susceptible tomatoes. We inoculated three *Solanum lycopersicum* tomato varieties: wilt-resistant Hawaii7996 (H7996), wilt-susceptible L390, and moderately wilt-susceptible Moneymaker (MM) via soil drench inoculation [20,49]. Alhough H7996 is wilt-resistant, *Ralstonia* is still able to colonize this variety, albeit at lower levels than wilt-susceptible plants [49,50]. *Rs* K60 colonization in roots was quantified at 24, 48 and 72 hours post infiltration (hpi). In all tomato varieties, the Δ*ripU^K60^* single mutant had significantly lower rates of colonization when compared to both wild-type *Rs* K60 and Δ*ripU::RipU^K60^* at all time points (∼10^4^ CFU/g root tissue compared to ∼10^7^ CFU/g root tissue for wild type or complemented strain, Figure 4A). Colonization rates of Δ*ripU ^K60^::RipU^K60^*mimicked those of WT *Rs* K60 and were not significantly different at any time point in any tomato variety (Figure 4A). These data indicate that a functional RipU^K60^ is required for full *Rs* K60 virulence in tomato roots.

**Figure 4.**
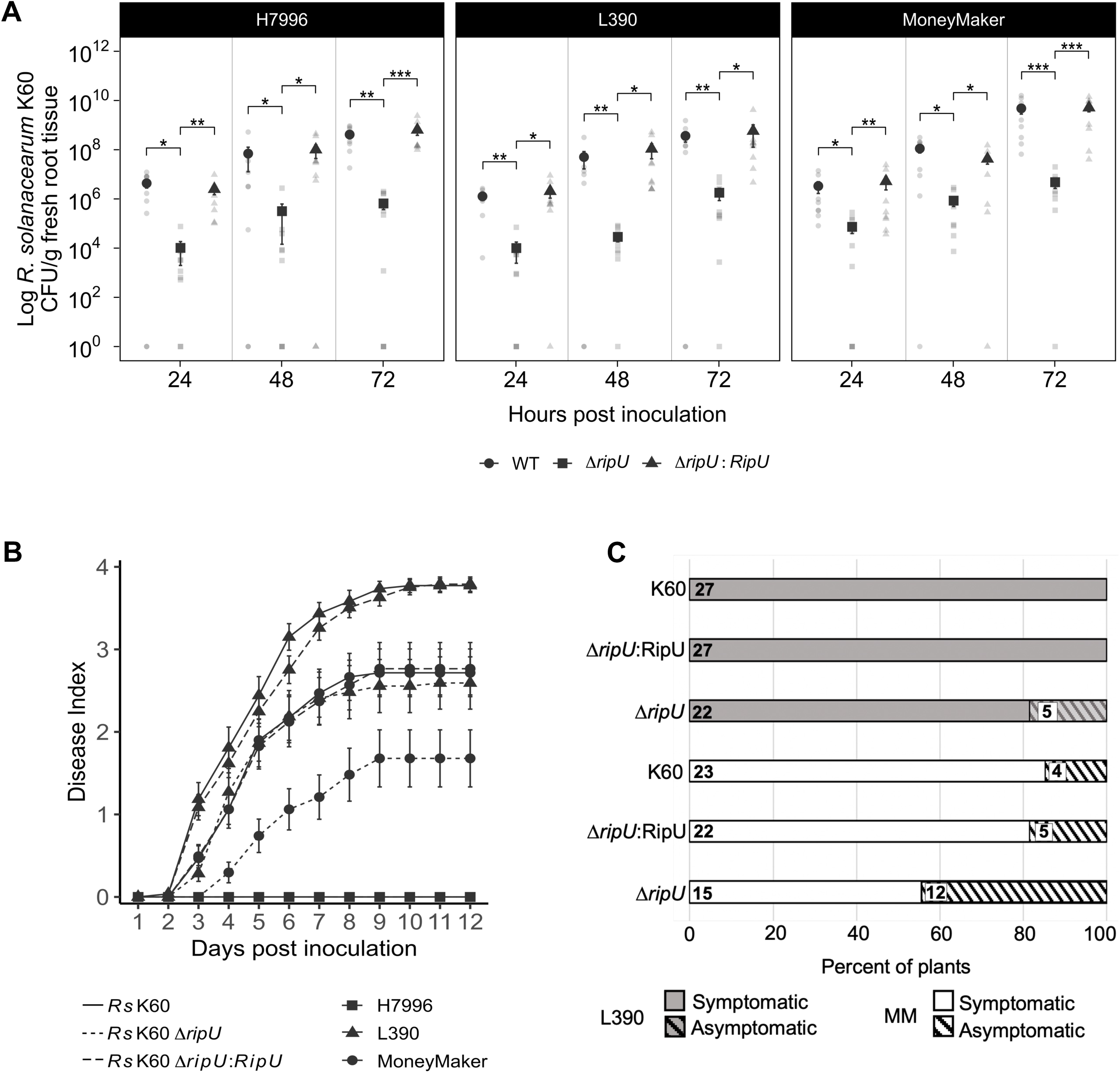
RipU^K60^ plays a role in *Rs* K60 pathogenesis and virulence. Root colonization of *Rs* K60, Δ*ripU^K60^* and Δ*ripU^K60^*::*RipU^K60^* in whole roots of H7996 (resistant), MM (susceptible) and L390 (susceptible) (A). Each dot represents one root (n = 9 per timepoint and genotype). Stars indicate significance with a Wilcox test (* = P < 0.05, ** = P < 0.01, *** = P < 0.001). Error bars indicate standard deviation. B and C. Infection with Δ*ripU^K60^*mutant results in less wilting symptoms in susceptible varieties. B. The Δ*ripU^K6^*^0^ mutant has delayed symptom development and less wilting at 12 dpi compared to either wild-type *Rs* K60 or Δ*ripU^K60^*:: *RipU^K60^*. Wilting was scored daily based on the number of leaves wilted per plant. Each point represents the average of 3 independent experiments, each with 12 plants per genotype per treatment. C. Fewer wilt-susceptible L390 and MM tomato plants show wilting symptoms when soil-drench inoculated with Δ*ripU^K60^*mutant strain compared to either wild-type *Rs* K60 or the complemented strain Δ*ripU^K60^*:: *RipU^K60^*. MM inoculated with any *Rs* strain reached the highest disease incidence at 7 dpi. L390 inoculated with wild type *Rs* K60 and the complemented strain reached highest disease incidence at 6 dpi.

Since the absence of RipU^K60^ influenced the ability of *Rs* K60 to colonize tomato roots, we next asked whether RipU^K60^ was required for *Rs* K60 pathogenicity. Wilt-resistant H7996 and wilt-susceptible MM and L390 plants were inoculated with either *Rs* K60, Δ*ripU^K60^*, or Δ*ripU^K60^::RipU^K60^*. Wilting was assayed on a scale of 0 to 4 (no discernable wilting = 0 and 100% of leaves wilted = 4). As expected, wilt-resistant H7996 plants inoculated with wild-type *Rs* K60, Δ*ripU^K60^*, or Δ*ripU ^K60^::RipU^K60^* did not display any observable wilting symptoms (score of 0 at 12 dpi; Figure 4B). Wilt-susceptible L390 plants inoculated with the Δ*ripU ^K60^* mutant displayed fewer wilting symptoms (wilting score of 2.59 at 12 days post inoculation (dpi)) than L390 plants inoculated with wild-type *Rs* K60 or Δ*ripU ^K60^::RipU^K60^*(score of 3.77 and 3.79 at 12 dpi, respectively; Figure 4B). Similarly, wilt-susceptible MM plants inoculated with the Δ*ripU^K60^*mutant displayed fewer wilting symptoms (score of 1.67 at 12 dpi; Figure 5B) than MM plants inoculated with wild-type *Rs* K60 or Δ*ripU ^K60^::RipU^K60^*(score = 2.71, and 2.67, respectively; Figure 4B). Additionally, the Δ*ripU^K60^* mutant had delayed virulence on susceptible tomato plants.

**Figure 5.**
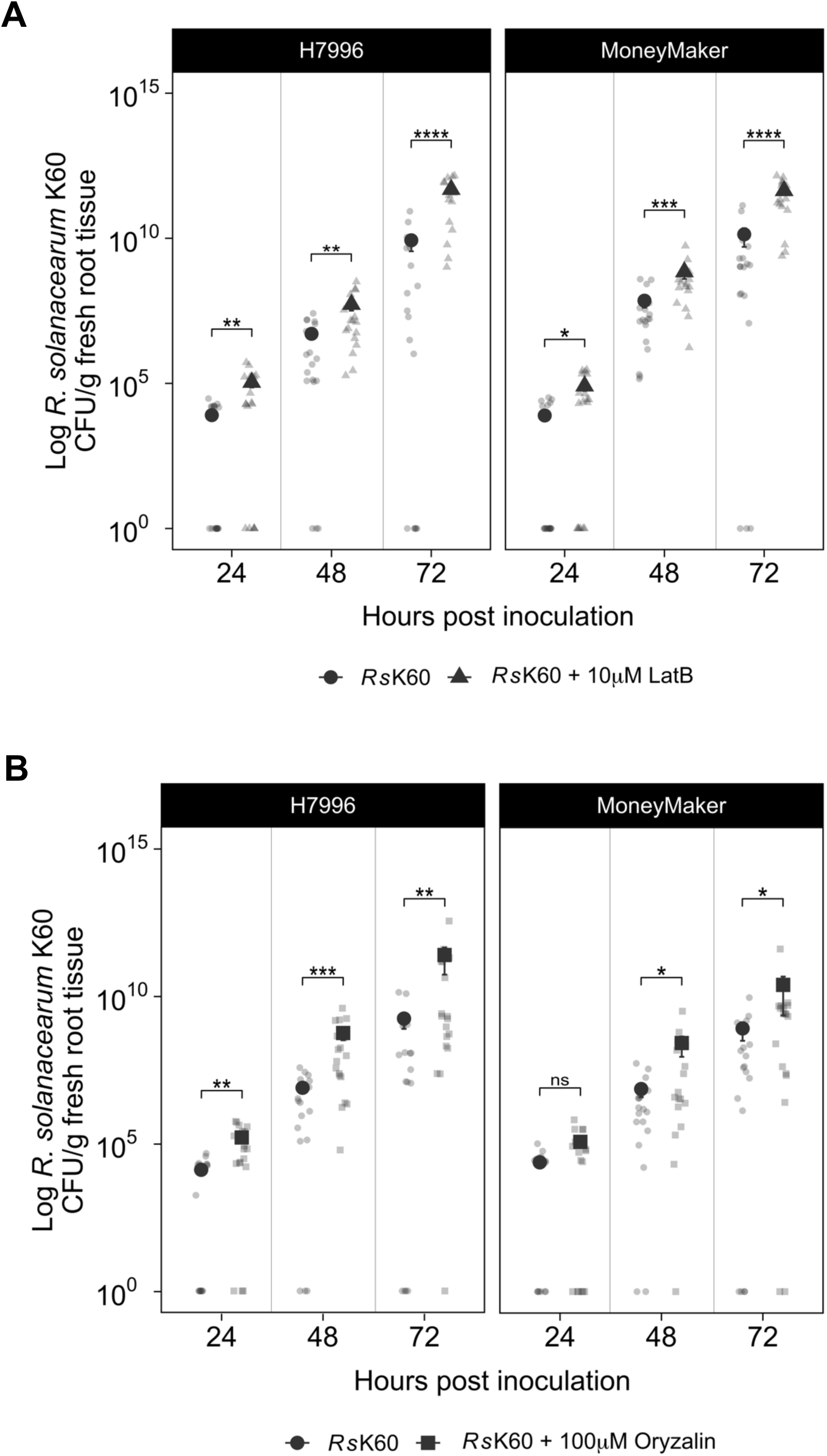
Cytoskeleton disruption improves root colonization in both resistant (H7996) and susceptible (Moneymaker) tomato plants. Tomato seedlings were grown on water agar and treated with 10µM latrunculin B (LatB) solution, 100µM oryzalin solution, or mock solution (0.5X MS + DMSO) two hours before inoculation with *Rs* K60. Graph A compares *Rs* K60 with *Rs* K60 LatB treatment and graph B shows *Rs* K60 compared to *Rs* K60 oryzalin treatment. 3 independent tests (6 samples per treatment per genotype). Wilcoxon Test: * = p<0.05, ** = p<0.01, ***= p<.005, ****= p<.001. Error bars = standard deviation.

Analysis of disease incidence (number of plants with any symptoms/number of total plants) showed that the disease incidence of both wilt-susceptible genotypes L390 and MM was lower when inoculated with the Δ*ripU^K60^* mutant compared to either wild type *Rs* K60 or the complemented *Rs* strain (Figure 4C). L390 plants treated with *Rs* K60 or the complemented Δ*ripU ^K60^::RipU^K60^* reached 100% disease incidence (27/27 plants) at 6 days after inoculation. In contrast, the highest disease incidence for L390 plants treated with the Δ*ripU^K60^* mutant was 81.4% (22/27 plants) at 8 dpi (Figure 4C). MM plants inoculated with any strain reached their highest disease incidence at 7 dpi. However, 85% (23/27) of MM treated with *Rs* K60 and 81.4% (22/27) of MM plants inoculated with the complemented strain showed wilting symptoms. In contrast, only 55.5% (15/27) of MM plants inoculated with the Δ*ripU^K60^* mutant had wilting symptoms (Figure 4C). These findings, taken together with data from the root colonization assays, demonstrate that RipU^K60^ is essential for full pathogenicity and virulence of *Rs* K60.

### Chemical disruption of the actin or microtubule cytoskeletons improves Rs K60 colonization

Our data indicated that RipU^K60^ is required for virulence and pathogenicity of *Rs* K60, and that this effector associates with both actin and microtubules. In addition to these findings, a meta- analysis of three RNAseq datasets from resistant and susceptible tomatoes infected with *Ralstonia* revealed that genes involved in cytoskeletal organization were enriched among downregulated genes in susceptible tomatoes, but not in resistant plants [51] (Supplemental Figure 3). Further, pharmacological disruption of the actin or microtubule cytoskeleton can increase bacterial colonization in other Arabidopsis-*P. syringae* interactions (Henty-Ridilla et al. 2013, Kang et al. 2014). Thus, we hypothesized that chemical disruption of the cytoskeleton would promote *Rs* K60 colonization. To test this, we treated roots of tomato seedlings with pharmacological inhibitors that disrupt microtubules or actin and inoculated with *Rs* K60. Latrunculin B (LatB) inhibits actin polymerization by binding to monomeric actin and preventing its assembly into filament ends [52]. Oryzalin promotes the depolymerization of microtubules [53]. Prior to *Rs* K60 inoculation, wilt-resistant and susceptible tomato seedling roots were pre-treated with 10 µM LatB, 100 µM oryzalin, or mock treatment solution (DMSO) for two hours. Roots were subsequently inoculated with *Rs* or water. Wilt-resistant Hawaii7996 (H7996) and wilt-susceptible Moneymaker (MM) plants pre-treated with LatB showed significantly increased *Rs* K60 colonization at 24, 48, and 72 hpi when compared to roots pre-treated with mock solution (Figure 5A). Thus, these data suggest actin disruption promoted *Rs* K60 colonization in both wilt-resistant and wilt-susceptible roots.

Similar to LatB, wilt-resistant H7996 pre-treated with oryzalin showed significantly increased *Rs* K60 colonization at 24, 48, and 72 hpi compared to roots pre-treated with mock solution (Figure 5B). In wilt-susceptible MM pre-treated with oryzalin, colonization of *Rs* K60 was significantly increased at 48 and 72 hpi (Figure 5B). Thus, pharmacological disruption of the actin or microtubule cytoskeleton promoted *Rs* K60 colonization in both wilt-resistant H7996 and wilt-susceptible MM roots.

### RipU^K60^ alters actin and microtubule organization

Given our findings that disruption of the cytoskeleton influences the ability of *Rs* K60 to colonize tomato roots, and that RipU^K60^ associates with the actin and microtubule cytoskeleton, we hypothesized that RipU^K60^ alters cytoskeleton organization to influence virulence. To investigate this hypothesis, we quantified actin and microtubule organization after transient expression of RipU^K60^ or RipBD^K60^ [19]. *N. benthamiana* leaves were co-infiltrated with *A. tumefaciens* strains transformed with either fABD2-mCherry or TUB5-mCherry, RipU^K60^-GFP or RipBD^K60^-GFP and single frame images collected from epidermal cells at 2 different timepoints with SDCM. Actin array organization was quantitatively assessed from images using parameters of percentage of occupancy or density as well as the Coefficient of Variation (CV) that describes the extent of filament bundling [54,55]. Transient expression of RipU^K60^-GFP did not significantly influence actin filament density or bundling compared to *N. benthamiana* leaves transiently expressing RipBD^K60^-GFP at 24 hpi. However, at 48 hpi actin filament density significantly increased while the extent of bundling significantly decreased in RipU^K60^-GFP treatments compared to the RipBD^K60^-control (Figure 6). Furthermore, *N. benthamiana* leaves transiently expressing RipU^K60^-GFP had significantly decreased microtubule density at 48 hpi (Figure 6C). Collectively, our results demonstrate that expression of RipU^K60^ disrupts cytoskeletal organization in *N. benthamiana*.

**Figure 6.**
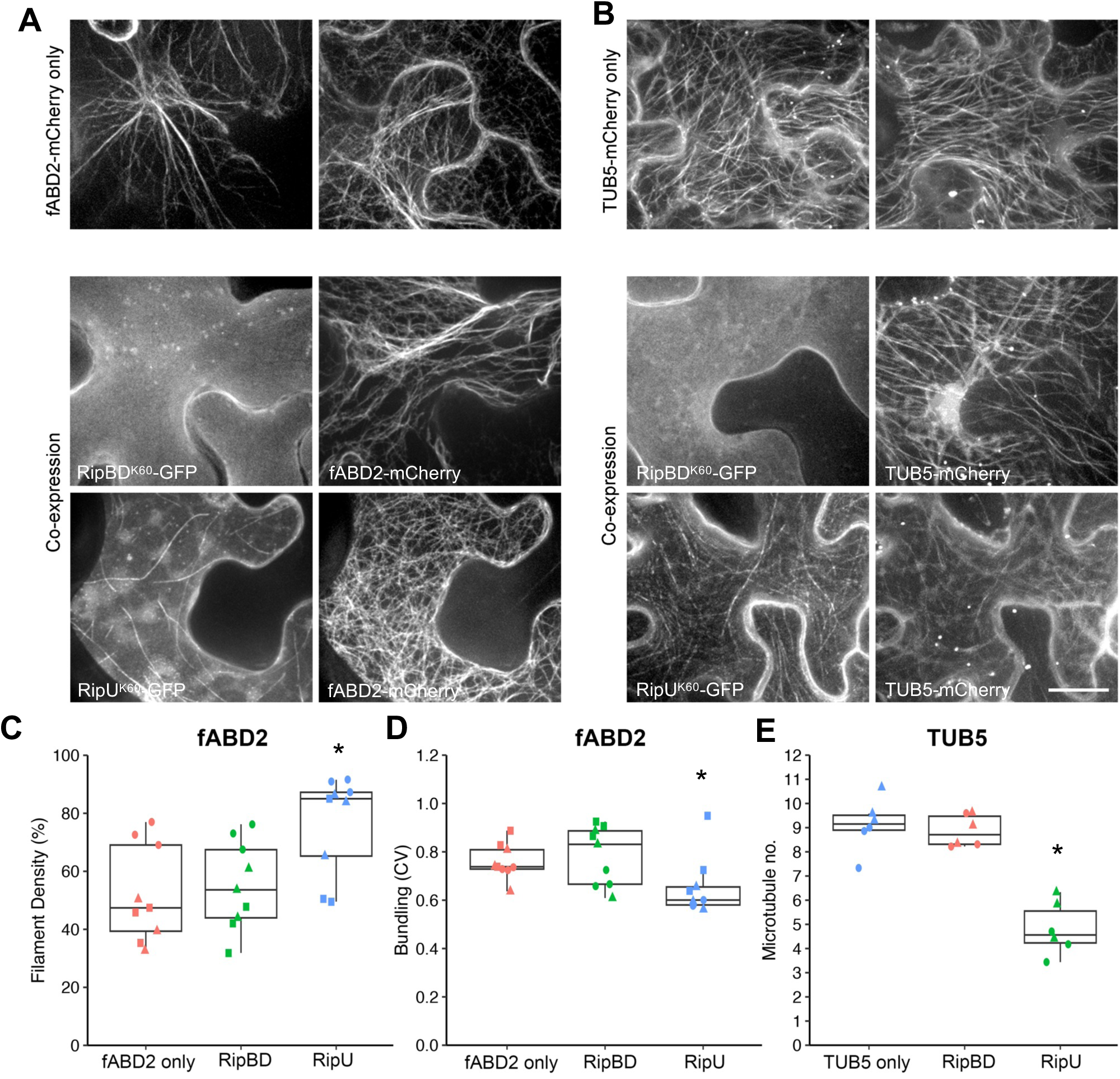
RipU^K60^ impacts plant cytoskeleton organization. RipU^K60^ or RipBD^K60^ were co- infiltrated with the either the actin marker fABD2-mCherry (A) or the β-tubulin marker TUB5- mCherry (B) in *N. benthamiana* leaves. As an additional control, cytoskeleton markers were infiltrated alone. Scale bar = 20 µm. Cytoskeleton dynamics were quantified at 48 hpi. RipU^K60^ significantly increased actin filament density (C), decreased actin bundling (D), and reduced microtubule density compared to controls (E). 5-15 cells were measured in each infiltration site and the values was averaged as one biological sample; n = 3 per biological repeat; One-way ANOVA; * P < 0.05.

### Cytoskeletal disruption restores Rs K60 ΔripU ^K60^ colonization

Having demonstrated that RipU^K60^ alters cytoskeleton organization and that cytoskeleton disruption improves *Rs* K60 colonization, we reasoned that pharmacological disruption of the cytoskeleton would restore the colonization ability of the Δ*ripU^K60^* mutant. To test this hypothesis, tomato seedlings grown on agar plates were pre-treated with 10 µM LatB, 100 µM oryzalin, or mock treatment solution for two hours until the treatment had soaked into the root and surrounding media. Roots were subsequently inoculated with either wild-type *Rs* K60 or the Δ*ripU^K60^* mutant. *Rs* colonization within the root tissue was examined at 24, 48, and 72 hpi. Inoculation with the Δ*ripU^K60^*mutant showed significantly reduced colonization in mock treated wilt-resistant H7996 and wilt-susceptible MM tomato roots at all three timepoints compared to wild-type *Rs* K60 (Figure 7). Significantly, there was no statistical difference in colonization patterns between tomato roots inoculated with *Rs* K60 or treated with a pharmacological cytoskeletal disruptor and then inoculated with Δ*ripU^K60^* (Figure 7). This finding was consistent across all three timepoints and both tomato genotypes. Our data thus support the conclusion that pharmacological disruption of both the actin and microtubule cytoskeleton rescues the Δ*ripU^K60^*colonization defect.

**Figure 7.**
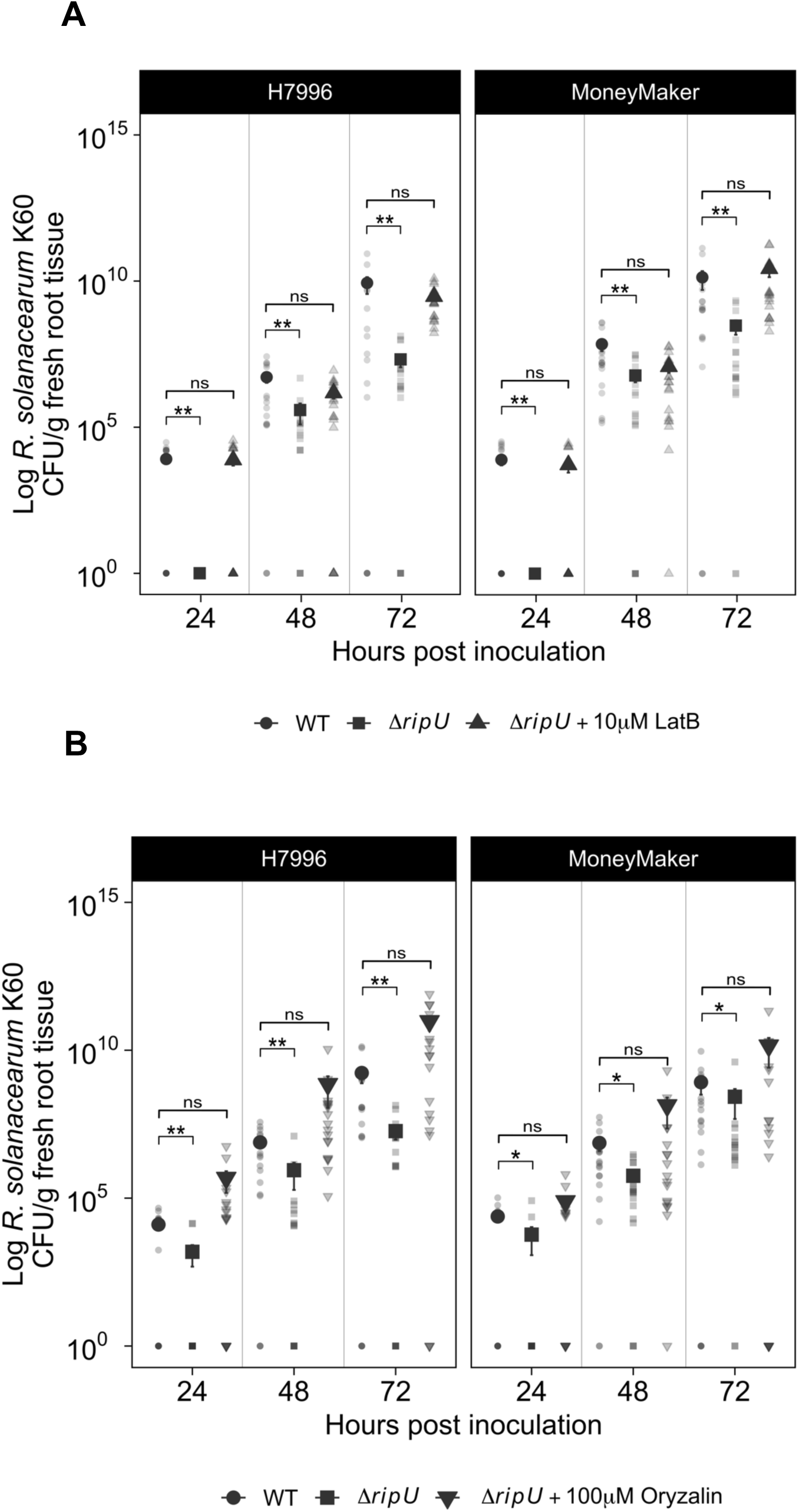
Chemicals that disrupt the cytoskeleton rescue the Δ*ripU* root colonization defect in wilt-resistant H7996 and wilt-susceptible Moneymaker. Tomato seedlings were grown on water agar and treated with (A) 10µM latrunculin B (LatB) solution, or (B) 100µM oryzalin solution, or mock solution (0.5X MS + DMSO) two hours before inoculation with either *Rs* K60 or Δ*ripU*. Three independent experiments were performed with six samples per treatment per genotype. Stars indicate significance with a Wilcoxon Test: * = P <0.05, ** = P <0.01. Error bars = standard deviation.

## Discussion

Here we show that a T3E protein from *Rs* K60 prominently colocalizes with microtubules, modestly colocalizes with actin filaments, physically associates with actin and tubulin, is required for bacterial virulence and colonization, and alters cytoskeleton organization in heterologous expression systems. We also show that an intact cytoskeleton functions in tomato immunity to *Rs*, as pharmacological disruption of either actin filaments or microtubules promotes *Rs* colonization in tomato roots. Further, we demonstrate that RipU is required for cytoskeleton-dependent proliferation within host plant tissues. Collectively, our data suggest that *Rs* K60 uses the T3E RipU^K60^ to promote its virulence *in planta* via an association with components of the cytoskeleton, likely interfering with cytoskeleton dynamics. We hypothesize that the changes in cytoskeleton organization and dynamics inhibit immune signaling, further enabling *Rs* virulence.

### Dynamic cytoskeletal rearrangements are a critical part of plant responses to microbes

Athough the cytoskeleton has been previously implicated in defense against other bacterial and fungal pathogens [4,5,7,41], our results represent significant advances in understanding the contributions of the host cytoskeleton in the tomato-*Rs* pathosystem. Actin remodeling is a broadly conserved plant immune response to cell surface-immune triggering microbes including the bacteria *Pseudomonas syringae, P. phaseolica*, *A. tumefaciens*, and the fungal pathogen *Magnaporthe grisae* [29]. For example, after inoculation with *P. syringae* pv *tomato* DC3000 a biphasic actin remodeling response is observed in which an initial increase in actin filament abundance occurs at 9 hpi, followed by enhanced bundling at 20 – 24 hpi [29]. Using different *P. syringae* strains and mutants, the initial response was found to be part of PTI and could be recapitulated with PAMPs, whereas the later response required the TTSS and effector proteins [29]. An increase in actin filament density is also observed in hypocotyl epidermal cells upon recognition of the immunogenic peptide elf26 [37,56]. Further, co-infiltration of Arabidopsis leaves with the actin polymerization inhibitor LatB and *P. syringae* pv *tomato* DC3000 leads to increased bacterial growth in leaves [29,30].

One of the best characterized actin remodeling events is the focal response to leaf penetration by fungi and oomycetes, in which cortical actin arrays are reorganized into a focal actin patch and radial bundles that are focused beneath the fungal contact site [32,33,35,42,57]. The reorganization of actin is accompanied by an increase in endoplasmic reticulum (ER) and Golgi bodies [32,42]. Actin-dependent transport enables polarized transport and secretion of cell wall and antimicrobial compounds directly to the infection site, resulting in cell wall barriers that slow pathogen invasion as well as a localized defense response [34,37,42,58–61]. In response to the powdery mildew pathogen *Blumeria graminis* f. sp. *hordei* (*Bgh*), the increased density of actin filaments at the infection site resembles an actin patch and precedes radial bundle accumulation [35]. Formation of the actin patch requires the actin nucleator proteins ARP2/3 and the class I formin AtFH1. The actin focal response is required to prevent fungal leaf penetration, as inhibiting actin rearrangements genetically or pharmacologically promotes pathogen penetration into host cells [35,62–65]

Although actin remodeling is part of innate immune signaling [29,36,37,56], the role of microtubules in innate immunity is less clear. This is partly because a range of microtubule organization changes are elicited by microbes and the types of changes depend on the genotype of both the plant and pathogen [41]. Arabidopsis susceptibility to the necrotrophic fungal pathogen *Sclerotinia sclerotiorum* was correlated with the ability of cortical microtubules to reorganize after infection [66]. Concentrations of microtubules form beneath fungal appressoria in barley (*Hordeum vulgare*) to prevent leaf penetration of *Blumeria graminis* f. sp. *hordei* [67,68] but also in flax mesophyll cells in response to an incompatible strain of the rust fungus *Melamspora lini* [69]. In contrast, microtubules reorient or depolymerize in resistant soybean cultivars in response to the oomycete *Phytophthora sojae* [70]. Chemical disruption of microtubules with oryzalin induces expression of defense genes in grapevine [71] and promotes susceptibility to virulent bacteria [31,46]. Microtubule reorganization can be important for fungal colonization, as inhibiting microtubule reorganization promotes penetration of non-host *Blumeria* in barley [63]. Unlike actin, changes in microtubule organization in Arabidopsis have not been observed in response to pathogenic bacteria or MAMPs [31,46], but have been observed in cells from other plant species such as *Vitis rupestris* (grapevine) cells [72]. While MAMP-elicited changes to microtubule dynamics remain poorly understood, changes in microtubule organization have been observed in response to other virulence factors such as microbe-produced toxins and proteins [73–75] and in response to beneficial microbes [76]. For example, high concentrations of *Verticillium dahliae* toxin (VD toxin) disrupt microtubules and reduce microtubule density in Arabidopsis [74,75]. The limited knowledge regarding the role of microtubules in immunity coupled with the complexity of these findings represent a significant knowledge gap that has yet to be addressed in cytoskeletal response to cell surface-immune signaling.

### What is the function of the RipU^K60^ induced cytoskeletal changes?

Transient expression of RipU^K60^ in *N. benthamiana* increased actin filament density and decreased bundling at 48 h following agroinfiltration. By preventing proper cytoskeleton remodeling, RipU^K60^ could repress cytoskeletal-mediated processes required for immune signaling, including immune receptor endocytosis and secretion of anti-microbial compounds, thereby enabling *Rs* colonization.

Cytoskeleton disruption may also indirectly promote *Rs* growth by enabling RipU^K60^ to gain better access to the vasculature. The microtubule cytoskeleton plays an important role in secondary cell wall formation in the xylem [77,78]. Trafficking of cellulose synthase complexes (CSCs) along cortical microtubules enables placement of cellulose in xylem cell walls [79–81]. As the xylem develops and undergoes programmed cell death, cell walls thicken and lignify. The denser, lignified cell walls pose a challenge for pathogen entry and are protective for the plant. Notably, not all areas of the xylem cell wall are lignified. Vascular pathogens like *Rs* can spread among xylem cells in part through pits which form between xylem cells [82]. Local depolymerization of microtubules prevents CSCs from moving to the region, thereby inhibiting secondary cell wall thickening [81,83]. By promoting the depolymerization of microtubules, RipU^K60^ may alter secondary cell wall structure, decreasing lignification and potentially increasing pit formation, thus enabling *Rs* more rapid entry into the vasculature.

### Elucidating the mechanisms underlying the interaction between RipU^K60^ and the cytoskeleton

We have shown that RipU^K60^ co-localizes and physically associates with components of the cytoskeleton. However, our data does not rule out that such association is direct or mediated by interactions with cytoskeleton associated proteins. Given that RipU^K60^ influences actin dynamics and microtubule number, RipU^K60^ may interact with components required for cytoskeleton remodeling, such as actin binding proteins (ABPs) and microtubule associated proteins (MAPs), or it may alter second messenger production (e.g. ROS, Ca^2+^, or phospholipids) that modulate cytoskeletal dynamics. Alternatively, RipU^K60^ could interfere with actin-microtubule crosstalk by associating with proteins that interact with both actin and microtubules [84]. The HopG1 effector from *P. syringae* triggers the actin reorganization observed during ETS [43] and interacts with a kinesin, a microtubule-associated motor protein. HopG1 co-immunoprecipitates with actin when kinesin is present but not when expressed alone [43]. Although the major function of kinesins is to move vesicles directionally along microtubules, these motor proteins can also cross-link actin to microtubules [85]. A kinesin mutant has reduced susceptibility to *Pst*, suggesting that HopG1 targets kinesin to promote actin changes that enhance pathogen virulence [43]. If RipU^K60^ interferes with actin-microtubule crosstalk, this may simultaneously alter both components of the cytoskeleton, actin immune signaling as well as downstream cell wall assembly resulting from microtubule disruption.

### RipU^K60^ is unusual among T3Es that impact the cytoskeleton

The cytoskeleton acts as a critical signaling intermediate during immune signaling, promotes immune receptor endocytosis and placement on the plasma membrane, and functions as a scaffold for vesicle trafficking and antimicrobial peptide transport [4,5,7]. Given its central role in immunity, it is perhaps unsurprising that microbial pathogens have evolved effector proteins that target and suppress cytoskeletal functions. Although multiple T3Es interact with the cytoskeleton, they do not alter its organization in the same way. For example, *Xanthomonas campestris* XopR directly interacts with the actin nucleating protein formin and promotes actin nucleation during early stages of infection. High concentrations of XopR inhibit nucleation and cause formin aggregation [39], although the relationship among XopR, the cytoskeleton, and symptom development is not clear. *Pst* DC3000 HopG1 induces actin bundling and decreases actin filament density in Arabidopsis cotyledons [43]. HopG1 does not impact *Pst* DC3000 growth in Arabidopsis but promotes symptom (chlorosis) development. The chlorosis induced by *Pst* DC3000 appears to be linked to changes in actin organization. Inoculating plants with DC3000 and inhibiting actin polymerization with cytochalasin D led to enhanced chlorosis, while promoting actin polymerization with jasplakinolide inhibited symptom development [43]. HopW1 from *Pst* co-immunoprecipitates with Arabidopsis Actin7 and, in contrast to RipU^K60^, reduces actin filament density when expressed in *N. benthamiana* or Arabidopsis at 6, 24 and 48 hpi [30]. HopW1 disrupts endocytosis during early infection in Arabidopsis [30] and may impact recycling of immune-related proteins at the cell surface. An effector with similarity to HopW1, the *Acidovorax citrulli* effector AopW1 group I, also disrupts the actin cytoskeleton, although the underlying mechanism is not completely understood [86].

Additional effectors target microtubules or microtubule-related proteins. HopZ1a, a T3E from *Pst* binds and acetylates tubulin [31]. Expression of HopZ1a causes a significant decrease in the density of microtubule networks in Arabidopsis, disrupts the plant secretory pathway, and suppresses callose formation [31]. Whether HopZ1a impacts the actin cytoskeleton is not known. HopE1 interacts with MAP65-1 in a calmodulin-dependent manner [46]. Binding of HopE1 from *Pst* DC3000 to calmodulin leads to the disassociation of MAP65-1 from microtubules but does not appear to impact the organization of microtubules. AvrBsT acetylates Arabidopsis ACIP1, which positively regulates immune responses and co-localizes with microtubules. Acetylation of AtACIP1 by AvrBsT changes this co-localization and promotes the aggregation of large AtACIP1 puncta throughout the cell. Whether microtubule localization is required for the immune-related function of AtACIP1 remains unclear.

To the best of our knowledge, only the T3E protein RipU^K60^ has been shown to physically associate with both actin and microtubules. However, several lines of evidence suggest that other effectors may also interfere with multiple structures of the plant cytoskeleton. Transient expression of XopL from *X. euvesicatoria* causes cell death in *N. benthamiama* and decreases microtubule number in *N. benthamiama* epidermal cells [45]. Co-localization of XopL with microtubules is correlated with the cell death phenotype, as XopL truncations that did not strongly co-localize with microtubules also did not cause cell death [45]. As discussed above, HopG1 alters actin organization but interacts with a microtubule motor protein. Although the impact of HopG1 expression on microtubule organization was not tested, these data suggest that interactions between actin and microtubules are important for host immune responses and suggest that proteins that interact with both actin and microtubules are possible targets for effector proteins.

### Conclusions

Effector proteins are often functionally redundant and/or work cooperatively to target common functions in host cells [1,18]. Targeting the cytoskeleton may serve not only as a platform for interfering with multiple plant functions but may provide a way for effector proteins to interact synergistically and gain additional functionality. Given the close relationship of the cortical cytoskeleton with the plasma membrane and cell wall [87], as well as the localization of many Rips to the cell periphery [19], we speculate that RipU^K60^ may function with other Rips to modulate immune processes at the cell periphery. For example, since microtubules influence cellulose alignment, one can hypothesize RipU functioning with another effector to alter cellulose synthase complex placement and cell wall structure. Alternatively, since actin promotes plasma membrane nanodomain formation [88], RipU^K60^ could function alongside other Rips to change the placement of immune receptors on the plasma membrane.

Together our data suggest that by preventing proper cytoskeleton remodeling, RipU^K60^ represses cytoskeletal-mediated processes required for immune signaling, thereby enabling *Rs* colonization. The discovery of a T3E that targets multiple components of the cytoskeleton underscores the importance of this proteinaceous network for immunity. The molecular mechanisms through which RipU^K60^ alters the cytoskeleton and the specific impacts of RipU^K60^- induced cytoskeletal disruptions (for example changes to PRR endocytosis) remain unknown but will be the subject of future investigation.

## Methods

### Plasmid construction

Full-length RipU^K60^ was PCR-amplified from *Rs* K60 genomic DNA and the resulting PCR products were cloned into either pENTR/D-TOPO or pBSDONR P1-P4 [89,90] Gateway donor plasmids using BP Clonase II (Invitrogen). We designated the resulting clones pENTR/D-TOPO- RipU^K60^ and pBSDONR (P1-P4)-RipU^K60^.

To generate the Green Fluorescent Protein (GFP)-tagged RipU^K60^ construct, the pENTR/D- TOPO-RipU^K60^ plasmid was mixed with the Gateway-compatible destination plasmid pB7FWG2, which places the transgene under the control of the 35S promoter [91]. Plasmids were recombined by addition of LR Clonase II (Invitrogen) following the manufactures instructions. The resulting expression clone was designated pB7FWG2-RipU^K60^-GFP.

To generate the pBAV154:RipU^K60^-3xHA expression construct, the pBSDONR (P1-P4)-RipU^K60^ plasmid was mixed with pBSDONR (P4r-P2):3xHA [89] and the Gateway-compatible destination plasmid, pBAV154, which places the transgene under the control of dexamethasone-inducible promoter [92]. The plasmids were recombined using LR Clonase II (Invitrogen) following the manufactures instructions. The resulting construct, pBAV154:RipU^K60^-3xHA, was used to generate transgenic Arabidopsis.

The pB7FWG2:RipU^K60^ and pBAV154:RipU^K60^-3xHA constructs were sequence-verified and subsequently mobilized into *Agrobacterium tumefaciens* GV3101 (pMP90).

### Agar plate-based plant growth conditions

Resistant tomato accession Hawaii7996 (H7996; *Solanum lycopersicum*), susceptible MoneyMaker (MM; *S. lycopersicum*) and susceptible L390 (*S. lycopersicum* var. *cerasiforme*) seeds were surface sterilized with a 50% bleach solution for 5 minutes, washed with water and stratified at 4°C overnight. Seeds were then planted on water-agar and grown at 28°CLto 30°C in a growth chamber at 16h:8h day/night cycle.

Arabidopsis (*Arabidopsis thaliana*) Columbia-0 (Col-0) plants were surface sterilized using 50% bleach and 70% ethanol and were plated on 0.5x Murashige and Skoog (MS) media containing nitrogen (Caisson Labs, UT, USA). Media was supplemented with various treatment solutions according to experiment. Seeds were stratified on treatment plates for 48 hours in the dark. Plates were then moved to a growth chamber at 22°C at 16h:8h day/night cycle.

### Soil-based plant growth conditions

Sterilized and stratified tomato seeds were planted in Pro-Mix PGX Propagation soil (Premier Tech Horticulture) in 3603 pots containing 25-27g of soil. Plants were grown under 16h:8h day/night cycle, at 28°CLto 30°C in a growth chamber for 15 days prior to inoculation with *Ralstonia*. At 10 days post planting, tomato seedlings were fertilized with Peters Fertilizer (Hummert International, USA).

*Nicotiana benthamiana* and *Nicotiana tabacum* seeds were sown in pots containing Pro-Mix PGX Propagation soil (Premier Tech Horticulture) and grown for 4 to 5 weeks in a growth chamber at 60% relative humidity in a 16-h-:8-h day/night cycle at 22°C to 23°C.

### Secretion assay

The secretion assays were performed as in [93]. The pAM5 plasmid (a gift from Stephane Genin) was electroporated into both *Ralstonia pseudosolanacearum* strain GMI1000 *(Rp*^GMI^) and Δ*hrcC^GMI^* mutant strains carrying RipU^K60^-3HA fusion. The Δ*hrcC^GMI^*mutant strain lacks a functional type III secretion system and is unable to secrete T3Es. The plasmid pAM5, which increases *HrpB* expression and allows for better detection of the effector in the culture supernatant, was electroporated into transformed *Rp^GMI^* or the Δ*hrcC^GMI^* mutant. Transformed *Rp^GMI^* and the Δ*hrcC^GMI^* mutant were cultured overnight at 28°C and pellets were resuspended in secretion media. After an 8h incubation at 28°C, samples were diluted with the secretion medium to ensure equal optical density (O.D.) readings (O.D._600_ 0.5-0.8). The supernatant and the pellet were then separated via centrifugation. Bacterial pellets were resuspended in sterile water and stored at -20°C. Supernatant samples were filtered, 1 mL of cold 25% trichloroacetic acid was added and incubated overnight at 4°C. For supernatant samples, the pellets were washed with 1 mL acetone 90%, the pellets dried, and stored at -20°C.

### Generation of transgenic Arabidopsis

The pBAV154:RipU^K60^-3xHA construct was transformed into wild-type Arabidopsis Col-0 using the floral dip method [94]. Progeny were selected for resistance to glufosinate in the T1 and T2 generations, and PCR amplification confirmed the presence of RipU^K60^ transgene.

### Agrobacterium-mediated transient expression in Nicotiana species

The pB7FWG2-RipU^K60^-GFP construct was transformed into *A. tumefaciens* GV3101 (pMP90) and streaked onto Luria-Bertani (LB) media supplemented with 25 μg gentamicin sulfate and 100 μg spectinomycin. Cultures were prepared in liquid LB, with the appropriate antibiotics, and grown overnight at 30°C. Following overnight incubation, cells were pelleted by centrifugation at 3,000 x g for 3 minutes at room temperature and resuspended in 10 mM MgCl_2_. Bacterial suspensions were diluted to an optical density at 600nm (OD_600_) of 0.5, incubated in 150 μM acetosyringone for 3-4 hours at room temperature, and infiltrated into four-week *Nicotiana benthamiana* or *N. tabacum.* For co-infiltration experiments, *A. tumefaciens* strain GV3101 harboring 35 S::fABD2-mCherry or UBQ10::TUB5-mCherry constructs were co-infiltrated into *N. benthamiana* with RipU^K60^-GFP or RipBD^K60^-GFP. Infiltrated leaves were imaged at 24 and 48 hpi.

### Cytoskeleton imaging and quantitative image analysis

*N. benthamiana* leaf epidermal cells co-expressing cytoskeletal markers and *Rs* T3Es were imaged by spinning disc confocal microscopy SDCM with an Olympus IX-83 inverted microscope equipped with a spinning disc confocal head (Yokagawa CSU-X1-A1; Hamamatsu Photonics, Hamamatsu, Japan) and an Andor iXon Ultra 897BV EMCCD camera (Andor Technology, Concord, MA, USA). Images were collected with an Olympus 60x oil objective (1.40 NA UPlanSApo; Olympus) using MetaMorph version 7.10.5 software. GFP and mCherry fluorescence were excited with 488-nm and 561-nm lasers and emission collected through 525/30-nm and 607/36-nm filters, respectively. Confocal z-series were taken at 0.5 μm step size for a total of 30 steps. For GFP and mCherry double-channel imaging, cells expressing single markers were checked to make sure no fluorescence bleed-through was detected in each channel. At least 5-10 images were taken at each infiltration site, and at least three infiltration sites were imaged for each treatment.

All image processing and analysis were performed in ImageJ or FIJI [95]. Epidermal cell z-series were converted into single images by maximum intensity projection before quantitative analysis. For colocalization analysis of RipU^K60^-GFP with actin or microtubule filaments, intracellular regions co-expressing both markers were cropped in both channels and used as Region of

Interests (ROIs) for Pearson’s correlation coefficient analysis. The analysis was performed with the ImageJ plug-in JaCoP [96]. Cells co-expressing RipBD^K60^-GFP and actin or microtubule markers were analyzed in the same experiments as control for random colocalization.

Actin density was analyzed as percentage of occupancy as previously described [29]. Actin extent of bundling was analyzed by quantifying the coefficient of variation (CV) of intensity values with an ImageJ macro developed by [55]. Microtubule density was estimated by counting microtubule numbers along a 10 µm line drawn vertically to the orientation of the most microtubules in a cell as previously described [97]. For all quantitative analysis, 5-15 cells were measured at each infiltration site and at least three infiltration sites were measured in each biological repeat.

### Luminol-based assay for quantification of Reactive Oxygen Species production in Nicotiana benthamiana leaves

This assay was performed as described previously [19,51]. *N. benthamiana* leaves infiltrated with *A. tumefaciens* were harvested at two days post infiltration. A 5 mm biopsy punch was used to generate leaf discs (four leaf discs per plant; 12 leaf discs in total for each construct). Discs were washed in water and kept in the dark for two hours, with water changes every 30 minutes. After two hours, leaf discs were transferred to a 96-well Perkin Elmer OptiPlate and stored overnight in water in the dark. The following day, leaf discs were treated with horseradish peroxidase and luminol solution, and a flagellin22 (flg22) peptide elicitor was added to induce ROS production. Three independent experimental replicates were performed. Area Under the Curve was calculated for each and compared using a Student T-Test in R version 3.6.1.

### Cell death suppression assay

Three-week-old *N. benthamiana* leaves were agroinfiltrated with *A. tumefaciens* GV3101 carrying pB7FWG2-RipU^K60^-GFP at an OD_600_ of 0.5, and infiltration sites were marked with a red permanent marker. Twenty-four hours post agroinfiltration (hpi), an overlapping region within the same *N. benthamiana* leaves were agroinfiltrated with *A. tumefaciens* GV3101 carrying the cell death elicitor, BAX, at an OD_600_ of 0.1, and the infiltrated regions were marked with a red permanent marker. Leaves were assessed for cell death five days after agroinfiltration with BAX and a representative leaf was photographed under white light.

### Callose deposition assay in Arabidopsis roots

The callose deposition assay was adapted from [98]. Wild type Columbia-0 (Col-0) and transgenic Arabidopsis plants with DEX-inducible RipU^K60^ expression were stratified for 48 hours and germinated on 0.5x MS media supplemented with 0.5% sucrose for seven days. Seedlings were then moved to 6-well tissue culture plates with liquid 0.5x MS (with sucrose) supplemented with pre-treatment solutions; DMSO, 10 µM DEX. Plants were grown in pre-treatment liquid media for three days, and media was replaced daily. Seedlings were then challenged with flagellin-22 (*Pseudomonas aeruginosa)* or a mock treatment solution for 24 hours. After MAMP treatment, seedlings were destained then transferred to a 0.01% aniline blue staining solution for two hours. Seedlings were plated in glycerol and imaged using a Zeiss 880 LSM Upright Confocal microscope (Carl Zeiss Micrscopy, White Plains, NY). Callose was imaged using the DAPI filter (358 nm excitation, 463 nm emission).

### Rs recombinant DNA techniques

A clean deletion mutant Δ*ripU^K60^* was created using *sacB* counter-selection with the vector pK18mobsacB as previously described [99]. Briefly, the upstream (646 bp) and downstream (536 bp) regions of *ripU* were amplified from *Rs K60* gDNA using Q5 high fidelity DNA polymerase (New England Biolabs, Ipswich, MA, USA) with the primers *ripU* up F/R and *ripU* dw F/R (Supplemental Table 1). Upstream and downstream fragments were fused and cloned at the HindIII site into pK18mobsacB by Gibson Assembly (New England Biolabs, Ipswich, MA, USA) following the manufacturer’s recommendations. pK18mobsacBΔ*ripU* construct was inserted into K60 using electroporation as previously described [100].

The first genomic recombination event was selected on CPG + Km. The second recombination event was screened for sucrose and Km sensitivity on CPG + 10% sucrose. Successful deletion of *ripU* (Δ659 bp) was confirmed using PCR with primers *ripU*flank F/R (Supplemental Table 1). Genomic DNA was isolated using Genomic DNA Buffer Set with Genomic-tip 20/G (Qiagen, Hilden, Germany). To construct the complementation vector, the gene region including the native promoter (405 bp upstream) and terminator (269 bp downstream) was polymerase chain reaction (PCR)–amplified from *K60 gDNA* and cloned via Gibson Assembly (New England Biolabs, Ipswich, MA, USA) at the HindIII site of pUC18miniTn7Gm to create pUC18miniTn7Gm::*ripU^K60^*, following the manufacturer’s protocol. K60Δ*ripU* was transformed with pUC18miniTn7Gm::*ripU^K60^* and pTNS3 to promote transposition and single gene insertion.

### Soil Drench Inoculation

Wild type *Rs* K60, the Δ*ripU ^K60^*mutant, and the Δ*ripU ^K60^::RipU ^K60^* complementation mutant were grown for two days on Casamino Peptone Agar (CPG) containing 1% triphenyl tetrazolium chloride (TZC) at 28^◦^C. Bacteria were harvested and resuspended in sterile water to 10^8^ CFU/mL. Tomato plants were inoculated at the three-leaf stage by applying inoculum (wild-type *Rs* K60, Δ*ripU ^K60^*, and Δ*ripU ^K60^::RipU ^K60^*) or water (mock treatment) directly to the soil using a serological pipette (1mL of inoculum/1 mg of soil) as described in Meline et al. 2022.

### Root colonization and disease index

To measure *Rs* K60 colonization, roots from inoculated tomato plants were harvested at 24, 48 and 72 hours post inoculation. Within treatments, harvested roots were pooled in groups of three and weighed after the removal of residual soil and water. The surface of these pooled roots was then sterilized. Root samples were ground with a mortar and pestle and resuspended with 1 mL of sterile water. This root tissue slurry was used to plate serial dilutions on CPG + 1% TZC to measure the colony forming units (CFUs) of *Rs* K60 present per gram of root tissue. These dilution plates were incubated at 28°C for 48 hours. To determine the pathogen titer, colonies were counted on the dilution plates and set relative to the mass of the original root tissue. Experiments were repeated in triplicate, with each experiment containing three biological samples per treatment, per time point. Data did not meet the assumption of normality; the Dunn Test was performed in RStudio version 3.6.1.

Inoculated tomato plants were scored daily for wilting severity to assess bacterial wilt disease index. Wilting symptoms were quantified on a scale from 0 to 4 (0 = no leaves with observable wilting, 4 = 100% of leaves wilting). Using these raw wilt scores, disease index was calculated using the following equation (*DI* = Disease Index, *n_w_* = number of leaves wilted, *n* = total number of leaves).

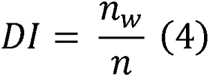

Experiments were replicated four times and each experiment consisted of eight to nine biological replicates.

### Cytoskeleton disruption and colonization assays

Tomato seedlings were grown on water agar and treated with either 10 μM latrunculin B (LatB) solution, 100 μM oryzalin solution, or mock treatment solution (0.5X MS + DMSO) two hours prior to inoculation with *Rs. Rs* was inoculated by pipetting 200 µl inoculum onto the entire root. Treatments were allowed to dry in a sterile hood. *Rs* K60, Δ*ripU^K60^* inoculum (1×10^5^ CFU/ml) or a mock treatment (water) was then applied to root tips. There were six treatments for these experiments; *Rs* K60, *Rs* K60 + LatB, *Rs* K60 + oryzalin, Δ*ripU^K60^*, Δ*ripU*^K60^ + LatB and Δ*ripU*^K60^ + oryzalin. Colonization assays were performed on inoculated roots. Colonization experiments were performed in triplicate, and each experiment consisted of three biological samples. Data were analyzed in R (version 3.6.1) using the Wilcox test.

### Co-immunoprecipitation assay

Co-immunoprecipitation (Co-IP) were conducted on protein extracted from *Nicotiana benthamiana* leaves expressing the green fluorescent protein (GFP) epitope tagged proteins as described previously [101] with slight modifications. Briefly, three leaves were harvested and pooled at 48hpi and preserved with liquid nitrogen. Leaf tissues were ground in 1mL of ice cold IP buffer (150 mM NaCl, 50 mM Tris [pH 7.5], 10% glycerol, 1 mM dithiothreitol, 1 mM EDTA, 1% Nonidet P-40, 0.1% Triton X-100, 1% plant protease inhibitor cocktail, and 1% 2,2’- dipyridyl disulfide) using a cold ceramic mortar pestle and were centrifuged at 10,000 x g for 15 minutes at 4°C. The supernatant was incubated with 10µL of green fluorescent protein (GFP)- Trap A (Chromotek) bead slurry for 4 hours at 4°C with constant slow rotation followed by washing the bead slurry 5 times with 500µL IP wash buffer at 1,000 x g for 1 minute at 4°C. The beads were resuspended in IP buffer and combined with 4x Laemmli buffer (BioRad) supplemented with 10% β-mercaptoethanol and the mixtures were boiled at 95°C for 10 min. 20 µl of input and 5 µl of IP samples were loaded and protein samples were separated on 4-20% Tris-glycine polyacrylamide gels (Bio-Rad) at 170 V for 1 hour in 1X Tris/glycine/SDS running buffer. Total proteins were transferred to nitrocellulose membranes (GE Water and Process Technologies) at 100 V for one hour. Membranes were incubated with blocking buffer (1X Tris-buffered saline (50 mM Tris-HCl, 150 mM NaCl [pH 6.8]) solution containing 0.1% Tween20 (TBST) containing 5% Difco skim milk) for 1 hour at room temperature with gentle shaking. Proteins were subsequently detected with horseradish peroxidase (HRP)-conjugated anti-GFP antibody (1:5,000) (Miltenyi Biotec #130-091-833), anti-plant actin mouse monoclonal antibody (3T3)-HRP (Abbkine # A01050HRP), alpha Tubulin monoclonal primary antibody (YL1/2) (1:10,000; Thermo Scientific MA1-80017) and goat anti-rat IgG secondary antibody (1:10,000; Thermo Scientific PA1-84709) in blocking buffer. Following antibody incubation, membranes were washed at least three times for 10 minutes with 1x TBST solution followed by 5 minutes incubation at room temperature with either Clarity Western ECL (BioRad). Immunoblots were developed using an Amersham ImageQuant 500 CCD imaging system (Cytiva). The experiment was repeated three times with similar results.

## Acknowledgements

This work was funded by a Foundation for Food and Agriculture (FFAR) New Innovator Award to AIP and the EMBRIO institute, contract #2120200, a National Science Foundation (NSF) Biology Integration Institute (CS and AIP). We are also grateful for support from the Ohio State University Presidential Student and Postdoctoral Fellowships to TLK and MVM, respectively. This research was also funded, in part, by the United States Department of Agriculture, Agricultural Research Service (USDA-ARS) research project 5020-21220-014-00D. The funding bodies had no role in designing the experiments, collecting the data, or writing the manuscript.

All opinions expressed in this paper are the authors’ and do not necessarily reflect the policies and views of USDA. USDA is an equal opportunity provider and employer. We thank Stephane Genin for the pAM5 plasmid, Katherine Rivera-Zuluaga for technical help with figure development and members of the Iyer-Pascuzzi and Jacobs labs for critical reading of the manuscript.

## Supplemental Information

**Supplemental Fig 1. RipU is secreted through the type III secretion system.** Immunoblots with RipU^K60^ tagged with HA detected in (A) both the supernatant and pellet of *Rp* GMI1000 and (B) only in the pellet of Δ*hrcC* mutant (RipU + HA tag = ∼38kDa).

**Supplemental Fig 2. RipU^K60^ fails to suppress BAX-triggered cell death in *N. benthamiana*.** Free GFP or RipU:GFP were transiently expressed in *N. benthamiana*. Twenty-four hours later the cell death-inducing construct, BAX, was infiltrated within an overlapping region of the initial infiltrated area. A representative leaf was photographed under white light 5 days following BAX infiltration. Black circles indicate areas infiltrated with either the GFP-tagged RipU or free GFP. Red circles represent areas infiltrated with the cell death-inducing construct.

**Supplemental Fig 3: Cytoskeleton-related Gene Ontology (GO) categories are enriched in a meta-analysis of *Ralstonia*-infected tomatoes.** (A) GO biological process categories related to the cytoskeleton that are enriched among downregulated genes in susceptible tomatoes in Meline et al. 2022. (B) Genes in the categories in (A) and their closest Arabidopsis homolog identified from Phytozome 13.

**Supplemental Table 1: Primers for *Rs* Recombinant DNA techniques**

