## Supplemental Figure 1 for "A *Ralstonia solanacearum* type III effector alters the actin and microtubule cytoskeleton to promote bacterial virulence in plants"

### Slide 1
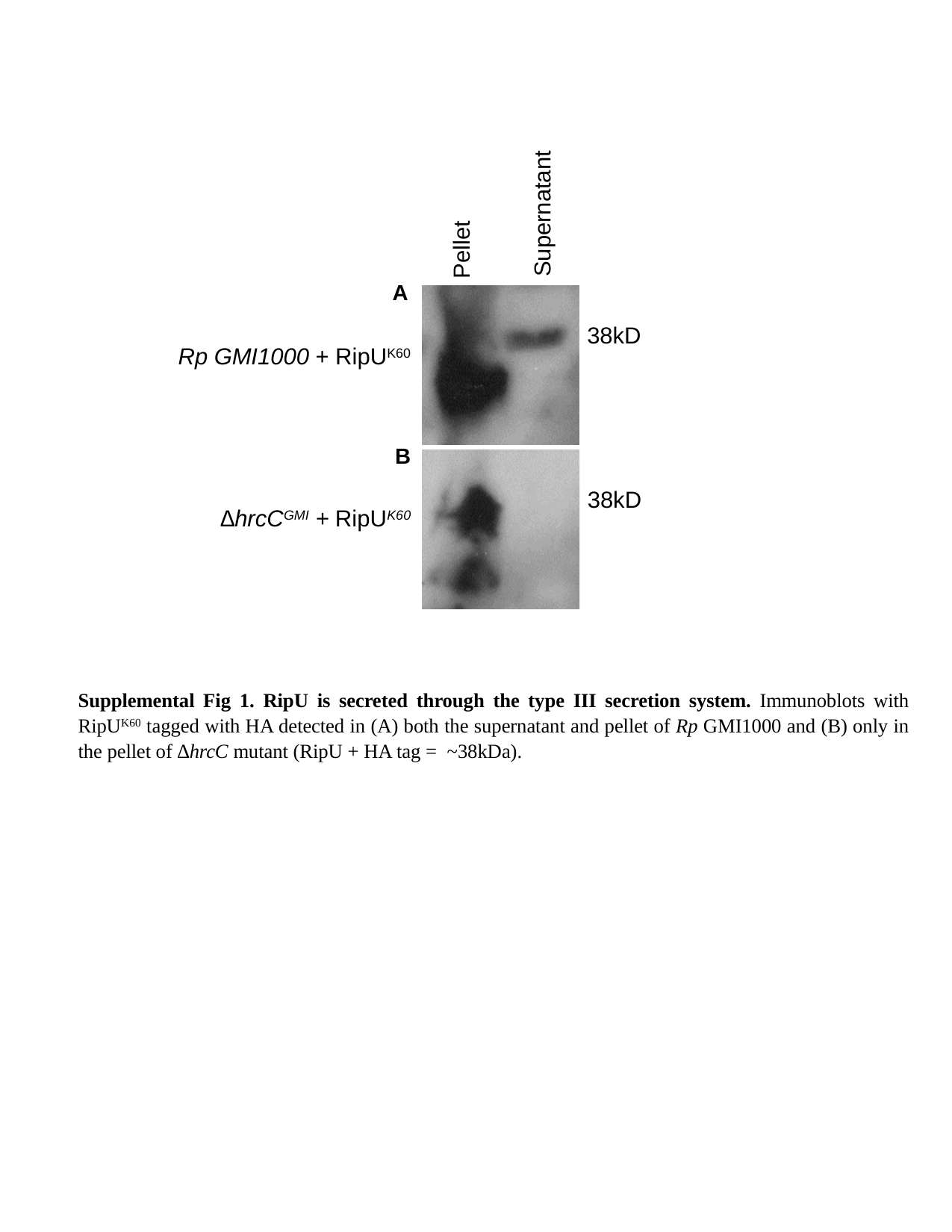

Supernatant
Pellet
38kD
Rp GMI1000 + RipU^K60^
38kD
∆hrcC^GMI^ + RipU^K60^
A
B
Supplemental Fig 1. RipU is secreted through the type III secretion system. Immunoblots with RipU^K60^ tagged with HA detected in (A) both the supernatant and pellet of Rp GMI1000 and (B) only in the pellet of ∆hrcC mutant (RipU + HA tag = ~38kDa).
