## Supplemental Figure 2 for "A *Ralstonia solanacearum* type III effector alters the actin and microtubule cytoskeleton to promote bacterial virulence in plants"

### Slide 1
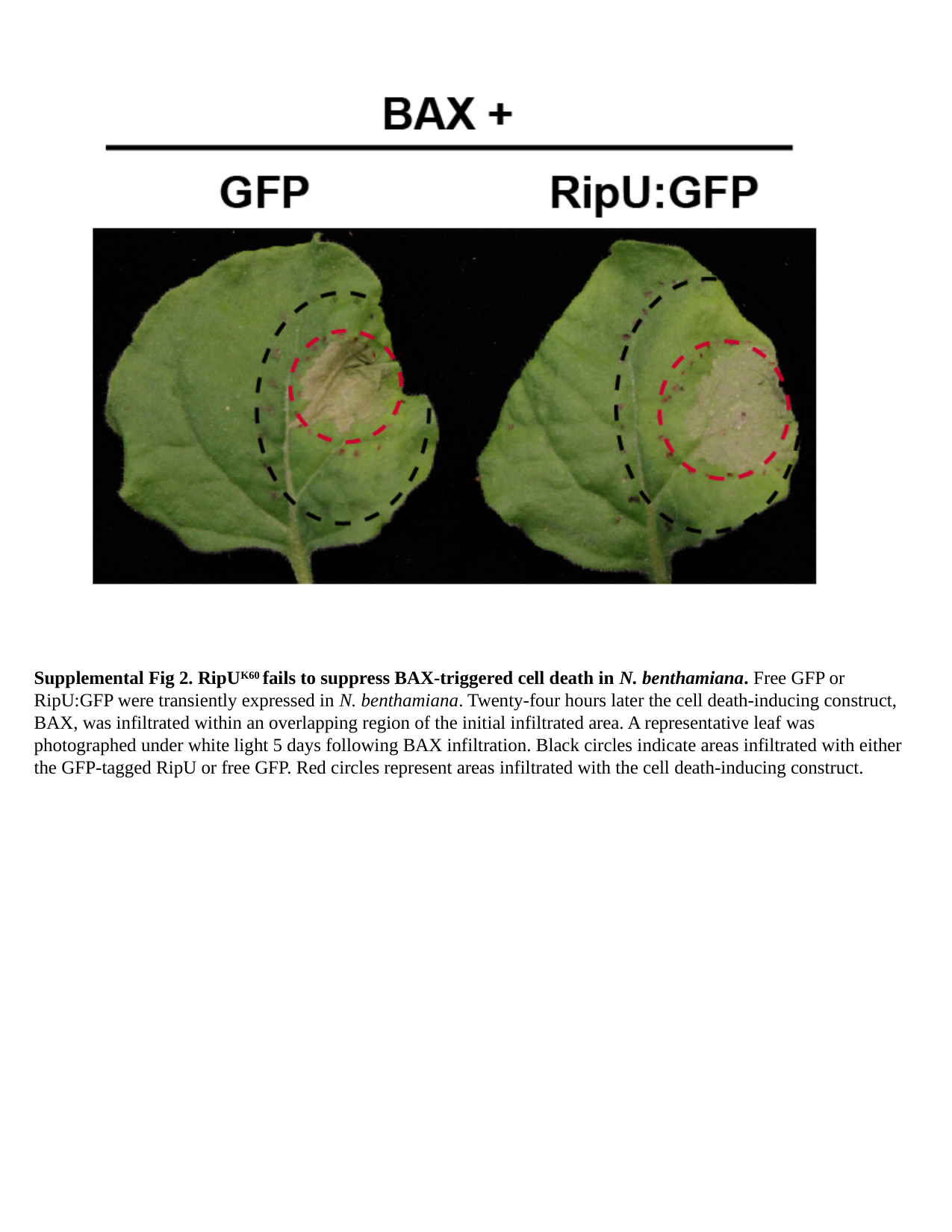

Supplemental Fig 2. RipU^K60^ fails to suppress BAX-triggered cell death in N. benthamiana. Free GFP or RipU:GFP were transiently expressed in N. benthamiana. Twenty-four hours later the cell death-inducing construct, BAX, was infiltrated within an overlapping region of the initial infiltrated area. A representative leaf was photographed under white light 5 days following BAX infiltration. Black circles indicate areas infiltrated with either the GFP-tagged RipU or free GFP. Red circles represent areas infiltrated with the cell death-inducing construct.
