## Supplemental Figure 3 for "A *Ralstonia solanacearum* type III effector alters the actin and microtubule cytoskeleton to promote bacterial virulence in plants"

### Slide 1
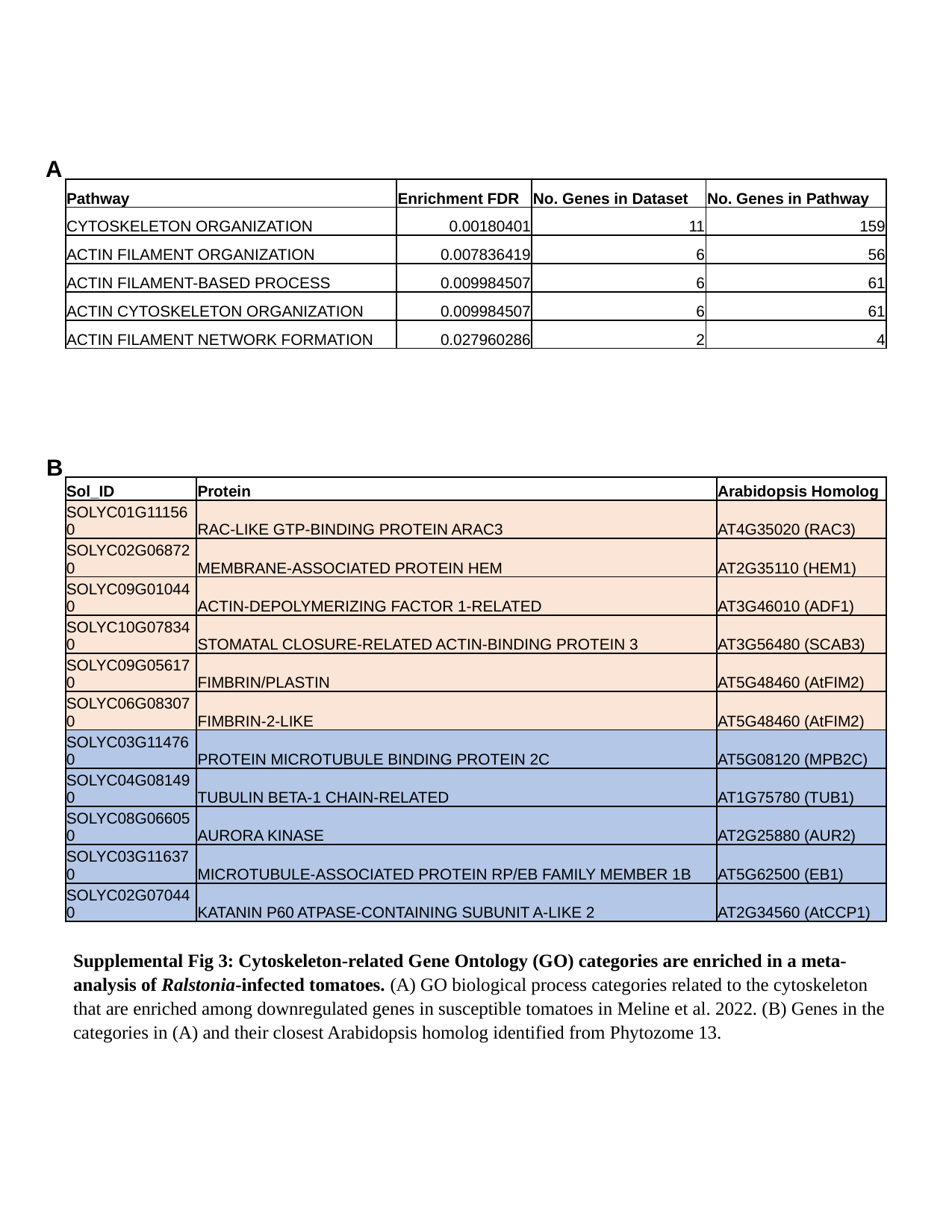

A
| Pathway | Enrichment FDR | No. Genes in Dataset | No. Genes in Pathway |
| --- | --- | --- | --- |
| CYTOSKELETON ORGANIZATION | 0.00180401 | 11 | 159 |
| ACTIN FILAMENT ORGANIZATION | 0.007836419 | 6 | 56 |
| ACTIN FILAMENT-BASED PROCESS | 0.009984507 | 6 | 61 |
| ACTIN CYTOSKELETON ORGANIZATION | 0.009984507 | 6 | 61 |
| ACTIN FILAMENT NETWORK FORMATION | 0.027960286 | 2 | 4 |
B
| Sol\_ID | Protein | Arabidopsis Homolog |
| --- | --- | --- |
| SOLYC01G111560 | RAC-LIKE GTP-BINDING PROTEIN ARAC3 | AT4G35020 (RAC3) |
| SOLYC02G068720 | MEMBRANE-ASSOCIATED PROTEIN HEM | AT2G35110 (HEM1) |
| SOLYC09G010440 | ACTIN-DEPOLYMERIZING FACTOR 1-RELATED | AT3G46010 (ADF1) |
| SOLYC10G078340 | STOMATAL CLOSURE-RELATED ACTIN-BINDING PROTEIN 3 | AT3G56480 (SCAB3) |
| SOLYC09G056170 | FIMBRIN/PLASTIN | AT5G48460 (AtFIM2) |
| SOLYC06G083070 | FIMBRIN-2-LIKE | AT5G48460 (AtFIM2) |
| SOLYC03G114760 | PROTEIN MICROTUBULE BINDING PROTEIN 2C | AT5G08120 (MPB2C) |
| SOLYC04G081490 | TUBULIN BETA-1 CHAIN-RELATED | AT1G75780 (TUB1) |
| SOLYC08G066050 | AURORA KINASE | AT2G25880 (AUR2) |
| SOLYC03G116370 | MICROTUBULE-ASSOCIATED PROTEIN RP/EB FAMILY MEMBER 1B | AT5G62500 (EB1) |
| SOLYC02G070440 | KATANIN P60 ATPASE-CONTAINING SUBUNIT A-LIKE 2 | AT2G34560 (AtCCP1) |
Supplemental Fig 3: Cytoskeleton-related Gene Ontology (GO) categories are enriched in a meta-analysis of Ralstonia-infected tomatoes. (A) GO biological process categories related to the cytoskeleton that are enriched among downregulated genes in susceptible tomatoes in Meline et al. 2022. (B) Genes in the categories in (A) and their closest Arabidopsis homolog identified from Phytozome 13.
